# Defining the HIV Capsid Binding Site of Nucleoporin 153

**DOI:** 10.1101/2022.05.06.490988

**Authors:** Shunji Li, Jagdish Suresh Patel, Jordan Yang, Angela Marie Crabtree, Brenda M. Rubenstein, Peik Karl Lund-Andersen, Frederick Marty Ytreberg, Paul Andrew Rowley

## Abstract

The interaction between the HIV-1 capsid (CA) and human nucleoporin 153 (NUP153) is vital for delivering the HIV-1 preintegration complex into the nucleus via the nuclear pore complex. The interaction with CA requires a phenylalanine/glycine-containing motif in the C-terminus of NUP153. This study used molecular modeling and biochemical assays to determine the amino acids of NUP153 that are essential for its interactions with CA. Molecular dynamics, FoldX, and PyRosetta simulations delineated the minimal CA binding motif of NUP153 based on the known structure of NUP153 bound to the HIV-1 CA hexamer. Computational predictions were experimentally validated by testing the interaction of NUP153 with CA using an *in vitro* binding assay and a cell-based TRIM-NUP153C restriction assay. This multidisciplinary approach identified eight amino acids from P1411 to G1418 that stably engage with CA, with significant correlations between molecular models and empirical experiments. Specifically, P1411, V1414, F1415, T1416, F1417, and G1418 were confirmed as critical amino acids required to interact NUP153 with CA.

**IMPORTANCE:** Human immunodeficiency virus (HIV) can infect non-dividing cells by interacting with host nuclear pores. The host nuclear pore protein NUP153 directly interacts with the HIV capsid to promote viral nuclear entry. This study used a multidisciplinary approach combining computational and experimental techniques to map the essential amino acids of NUP153 required for HIV capsid interaction. This approach revealed that the HIV capsid interacts specifically with only six amino acids of NUP153, suggesting other FG-containing motifs could also interact with the capsid. Based on molecular modeling, naturally occurring polymorphisms in human and non-human primates would be predicted to prevent NUP153 interaction with capsid, potentially protecting from HIV infection.

## Introduction

In eukaryotic cells, the nuclear envelope compartmentalizes the cytoplasm from the nucleoplasm. It is a physical barrier that must be traversed by viruses requiring access to the nucleus during their lifecycle, particularly when infecting non-dividing cells (1, 2). Access to the nucleus is through membranous pores in the nuclear envelope that are each stabilized by a large assemblage of ∼30 different nucleoporin proteins called the nuclear pore complex (NPC) (3). The NPC regulates nucleocytoplasmic transport with a selectively permeable barrier of unstructured filamentous nucleoporins that fill the nuclear pore and project from its surface. These filamentous nucleoporins contain an abundance of phenylalanine/glycine (FG) repeats that create a hydrophobic barrier to prevent the free diffusion of large macromolecules. Cellular proteins interact directly with nucleoporins to enable the nuclear ingress and egress of specific cellular cargos and interaction with FG nucleoporins is important for the efficient trafficking of macromolecules that are larger than ∼40 kDa.

Lentiviruses require the NPC to transport viral proteins and nucleic acids during the infection of non-dividing cells. Specifically, the HIV-1 genome is delivered to the NPC encapsidated in a fullerene cone constructed of capsid (CA) hexamers and pentamers (4, 5). The CA multimer has been observed docking with the surface of the NPC, but there is currently much debate on the exact mechanism of HIV-1 nuclear ingress (6). Many HIV-1 proteins have been shown to traffic to the nucleus, but the CA plays a dominant role in enabling the infection of non-dividing cells (7, 8). Genome-wide RNA interference screens have identified several nucleoporins required to complete the HIV-1 lifecycle (9–12). Of all the nucleoporins depleted from human cells in large-scale screens, NUP153 was consistently important for HIV-1 infection. NUP153 depletion resulted in up to a 100-fold drop in HIV-1 infectivity, reduced nuclear import of cDNA, and integration (13–17). The importance of NUP153 during viral nuclear ingress extends to other primate lentiviruses (18). However, it is less critical for lentiviruses that infect other mammals, such as equine infectious anemia virus and feline immunodeficiency virus (16).

NUP153 has an overall disordered structure and is anchored by its N-terminal domain to the nuclear basket. The NUP153 C-terminal domain (NUP153C) is rich in FG motifs that can project into the cytoplasm and nucleoplasm (19). The FG-rich C-terminus of NUP153 is required for CA binding (20), with a motif at amino acid positions 1407-1422 (TNNSPSGVFTFGANSS) playing a dominant role in this interaction (Figure 1A) (16, 21). NUP153C interaction does not occur with CA monomers and is specific to a hydrophobic pocket at the interface between adjacent monomers of the CA hexamer (21). Mutations within CA that disrupt this pocket can prevent HIV-1 infection of non-dividing cells and nuclear ingress of the preintegration complex. The central FTFG sequence of this motif (amino acids 1415-1418) is crucial for the interaction with a hydrophobic pocket formed between CA monomers. Moreover, mutation of the amino acids F1415, T1416, and F1417 in NUP153 interferes with the CA interaction (16, 21). Similarly, other host proteins also interact with CA at the same interface as NUP153, including CPSF6 and SEC24C (22, 23). All these proteins insert phenylalanine sidechains into the same hydrophobic pocket but have differences in the surrounding amino acid sequence. CA-targeting small molecules PF74, BI-2, and Lenacapavir also insert phenyl groups into the same binding pocket (24–26). Although there is an abundance of phenylalanine residues in many nucleoporins, there is a specific interaction of the NUP153 motif 1407-1422 with CA (16, 21). However, the sequence-specific determinants of this interaction have not been rigorously identified.

**Figure 1.**
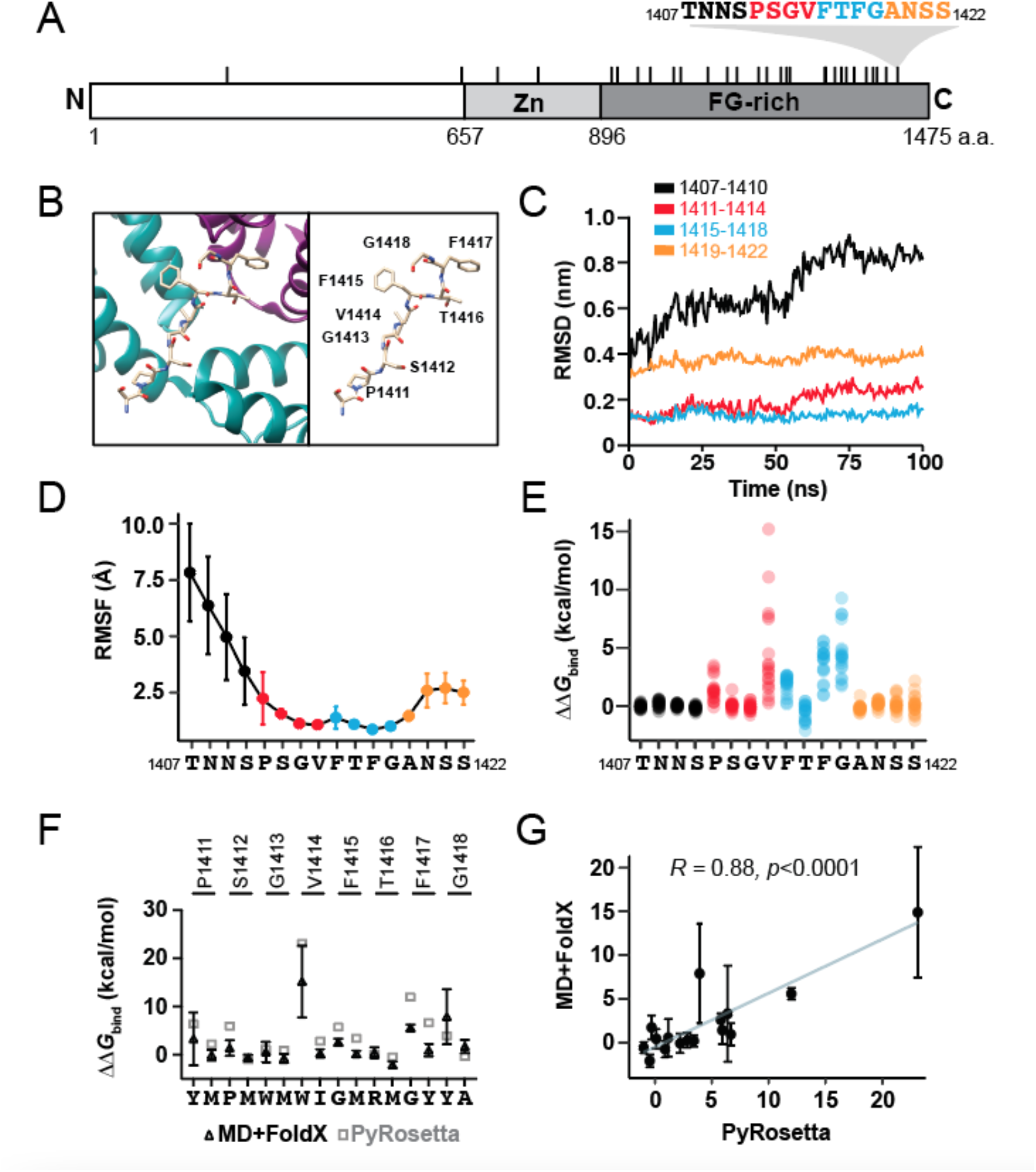
Molecular modeling of NUP153C-CA interaction defines a central eight amino acids that are stably associated with CA. (A) Domain diagram of the NUP153 protein. Tick marks represent FG repeats, with an expanded view of the FG motif that interacts directly with the HIV-1 CA (colored in relation to panels C-E). The numbering of NUP153 represents amino acid residues at the termini and domain boundaries. Zn -zinc finger domain. (B) *Left:* A structural representation of the HIV-1 CA hexamer bound by the NUP153C peptide (PDB: 4U0C). Magenta: monomer A; Green: monomer B. *Right:* A labeled representation of NUP153C without the CA structure. (C) RMSD of different regions of the NUP153C peptide during a 100 ns MD simulation. (D) A plot of the average RMSF of each amino acid residue in six copies of the NUP153C residues 1407-1422 bound to the hexamer CA during a 100 ns MD simulation (n= 6, error bars are standard deviations). (E) ΔΔ*G*_bind_ calculated by the MD+FoldX approach for all possible amino acid substitutions at each position in NUP153C (residues 1407-1422). (F) A comparison of the ΔΔ*G*_bind_ values calculated by MD+FoldX and PyRosetta for 16 NUP153C mutations. Error bars are standard deviations. (G) Correlation plot of ΔΔ*G*_bind_ estimated by MD+FoldX and PyRosetta, where the trendline shows the linear relationship between predicted ΔΔ*G*_bind_ values from the two different methods. Corresponding *R* and *p* values are displayed.

This study aimed to use molecular modeling combined with laboratory techniques to characterize the known interaction between NUP153 and HIV-1 CA. Molecular modeling has been utilized in many biological systems to answer fundamental questions regarding protein folding and function (27–29) and provides detailed information about how protein residues interact with a binding partner at the atomic scale. There are 29 FG repeats within NUP153, and it is unclear what makes the motif at positions 1407-1422 unique in its specific interaction with CA. Molecular dynamics (MD) simulations and *in silico* mutagenesis were used to determine the residues required for CA interaction with NUP153. These modeling predictions were validated by assaying mutant NUP153 and its interaction with CA in cell-based and *in vitro* CA pulldown assays. We find that modeling predictions correlate well with empirical studies. The stable interaction of specific residues of NUP153 with CA and their sensitivity to mutation has enabled the determination of the specific sequence motif within NUP153 required for interaction with CA.

## Results

### Molecular modeling identified residues essential for CA-binding in NUP153

The FG-containing motif of NUP153 interacts with a hydrophobic pocket formed by two adjacent CA monomers (Figure 1B) (21). The first molecular modeling approach involved conformational sampling of this CA-NUP153C complex via molecular dynamics (MD) simulations (30). Employing MD allowed us to investigate subtle conformational changes during the simulation and provided information on the stability of the interaction of NUP153C with CA. The root mean square deviation (RMSD) calculated using the NUP153 peptide backbone atoms confirmed that amino acids P1411-G1418 were stably associated with CA during 100 ns MD simulations (Figure 1C). Root mean square fluctuations (RMSF) analysis indicated that amino acids P1411-G1418 fluctuated less while in the binding pocket of CA during MD simulation (RMSF < 2.5 Å) (Figure 1D). Larger RMSF values of amino acids 1407-1410 and 1419-1422 indicated that they did not stably interact with CA during these simulations. The protein-protein binding affinity prediction tools FoldX and PyRosetta were then employed to assess the effects of amino acid substitutions in NUP153C on the binding stability using the ΔΔ*G*_bind_ value, where ΔΔ*G*_bind_ = Δ*G*_bind_ (mutant) - Δ*G*_bind_ (wild-type). The FoldX analysis combined the FoldX software with MD simulations to compute ΔΔ*G*_bind_ values for all possible 19 amino acid substitutions at each site in the NUP153 motif (27). Overall, a negative ΔΔ*G* value suggests the binding is stabilized by an amino acid substitution, whereas a positive value indicates destabilization. Five amino acid residues, P1411, V1414, F1415, F1417, and G1418, were considered critical binding sites for the CA interaction, as substituting these residues resulted in more positive ΔΔ*G*_bind_ values (Figure 1E). These residues are part of the region in NUP153C (P1411-G1418) that is stably associated with CA during MD simulations. The ΔΔ*G*_bind_ values were compared to each substitution’s volume and hydrophobicity values (Table 1). Increased CA binding (decreasing ΔΔ*G*_bind_ values) was significantly correlated with the mutations that altered the volumes of the amino acid sidechains. For the residues F1415-F1417, increasing sidechain volume increased binding, whereas smaller sidechain volume at residues G1413, V1414, and G1418 increased binding (Table 1). CA binding at position T1416 was correlated with increased sidechain volume and hydrophobicity (Table 1).

**Table 1.** The volume of amino acid sidechains in NUP153C is more important for CA interaction than hydrophobicity. Pearson correlation coefficient values comparing the ΔΔG_bind_ MD+FoldX against the hydrophobicity and volume of amino acid sidechains. * *p* <0.05, ** *p* <0.01.

| Residue | Pearson correlation coefficient |  |
| --- | --- | --- |
|  | Hydrophobicity | Volume |
| P1411 | -0.368 | 0.4068 |
| S1412 | -0.4023 | -0.4348 |
| G1413 | -0.4193 | 0.4738* |
| V1414 | -0.2894 | 0.6345** |
| F1415 | -0.3076 | -0.7852** |
| T1416 | -0.6186** | -0.4828* |
| F1417 | -0.4132 | -0.7489** |
| G1418 | -0.2615 | 0.7582** |

To validate the predictions made by MD+FoldX, we selected two substitutions at each position from P1411-G1418 for analysis with PyRosetta. These 16 mutations represented substitutions with high or low ΔΔ*G*_bind_ values. Overall, PyRosetta agreed with the MD+FoldX predictions for mutations at positions G1413, T1416, and G1418 and predicted larger ΔΔ*G*_bind_ values for the destabilizing mutants of P1411, S1412, V1414, F1415, and F1417 (Figure 1F). Comparing the predictions of MD+FoldX and PyRosetta resulted in a strong positive correlation (Pearson’s correlation coefficient r = 0.88, *p* <0.0001) (Figure 1G).

### Molecular modeling predicts the effects of mutations in NUP153C on the CA interaction as measured by co-sedimentation

Molecular modeling predictions suggest that specific amino acids were more important for the NUP153C interaction with CA. To validate modeling predictions from both MD+FoldX and PyRosetta, 16 mutations in the central PSGVFTFG motif (residues 1411-1418) were created in NUP153C to represent eight substitutions with high ΔΔ*G*_bind_ and eight with low ΔΔ*G*_bind_. Each mutant NUP153C was expressed in a HEK293T human cell line as a TRIM domain fusion and a C-terminal HA tag. Cell lysates containing NUP153C were used to determine interaction with recombinant purified multimeric CA tubes (Figures 2A and 2B). To assemble these tubes, CA monomers with engineered cysteine mutations were cross-linked to form hexamers (Figure 2A) and assembled into higher-order tubular structures (Figure 2B). The NUP153C-CA interaction was determined based on the fraction of NUP153C that bound and co-sedimented with multimeric CA tubes (Figure 2C). Wild-type NUP153C was efficiently bound to CA tubes, with 46% detected in the pellet fraction (Figure 2C and 2D). The binding of NUP153C with CA depended on the formation of the multimeric CA tubes as the reduction of the cystine bonds disassembled the CA tubes and localized NUP153C to the supernatant fraction (Figure 2C). A mutant NUP153C with a deletion of the entire interaction motif (ΔP1411-G1418) resulted in only 26% of NUP153C binding to CA (Figure 2D). As predicted by molecular modeling, the substitutions with low ΔΔG_bind_ (S1412M, G1413M, F1415M, F1417Y, and G1418A) did not reduce the binding with CA compared to wild-type NUP153C (Figure 2D). The mutations G1413M, F1415M, and F1417Y significantly increased the binding of NUP153C to CA tubes compared to the wild-type. Destabilizing mutations that had the highest ΔΔ*G*_bind_ values (average of 4.62 ± 4.98 kcal/mol MD+FoldX, 7.295 ± 7.349 kcal/mol PyRosetta) displayed reduced binding to CA tubes similar to the deletion mutant (ΔP1411-G1418) (Figure 2D). The mutants S1412P and G1413W appeared to bind CA similar to wild type NUP153C, reflecting their lower ΔΔ*G*_bind_ values (1.413/0.544 kcal/mol MD+FoldX and 5.919/1.159 kcal/mol PyRosetta, respectively). The conservative mutations P1411M, V1414I, and T1416M did not bind with CA regardless of their lower ΔΔ*G*_bind_ values (Figure 2D).

**Figure 2.**
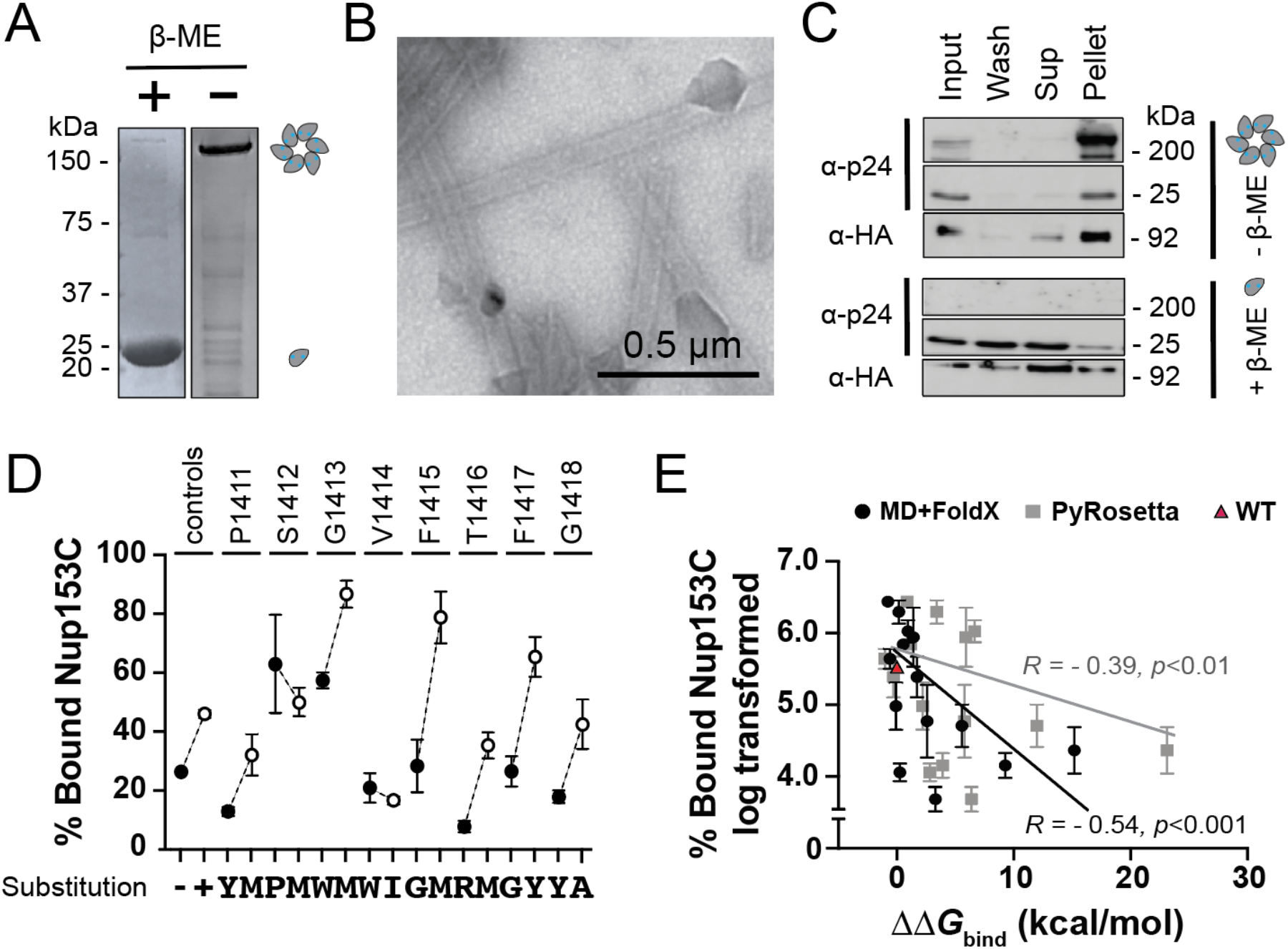
Molecular modeling predicts the effects of NUP153C mutations on CA interaction as measured by co-sedimentation. (A) SDS-PAGE of the monomeric and the cross-linked hexameric CA. (B) Transmission electron micrograph of CA tubes assembled from cross-linked CA hexamers. (C) A Western blot of a co-sedimentation assay using CA monomers (with the reducing agent β-mercaptoethanol (β-ME)) and CA tubes (without β-ME). Input: 10% of the total reaction volume. NUP153C was detected with an anti-HA antibody. (D) NUP153C mutants co-sedimented with CA tubes. Western blot signals were normalized to input. Error bars are standard deviations. Control reactions included the wild-type NUP153C (+) and NUP153C with a deletion of the CA interaction motif Δ1411-1418 (-). White data points represent mutations with low ΔΔ*G*_bind_ values, and black circles represent mutations with high ΔΔ*G*_bind_ values for each site. (E) Evaluation of the ΔΔ*G*_bind_ calculated by either MD+FoldX (black) or PyRosetta (gray) versus the experimental dataset. The Pearson’s correlation was calculated for each modeling method against three independent replicates of experimental data.

These results suggested that P1411, V1414, F1415, T1416, F1417, and G1418 are necessary for CA interaction as they appeared most sensitive to mutation. S1412 or G1413 were less critical for CA interaction and could tolerate mutations predicted to increase ΔΔ*G*_bind_. A comparison of these binding data with modeling predictions shows a significant negative correlation (Pearson’s correlation coefficient -0.54 (*p*<0.001) and -0.39 (*p*<0.01), for MD+FoldX and PyRosetta, respectively) (Figure 1E).

### Mutations predicted to disrupt NUP153C-CA interaction prevent HIV-1 restriction by TRIM-NUP153C

Modeling predictions made by MD+FoldX and PyRosetta were further scrutinized by testing the interaction between NUP153C and CA using a cell-based assay (Figure 3A) (16). HEK293T cells were transiently transfected with NUP153C with an N-terminal fusion to the TRIM domain from the Rhesus Macaque TRIM5α restriction factor. Cell lines expressing TRIM-NUP153C were challenged with HIV-GFP pseudotyped with VSV-G (Figure 3A). Interaction between TRIM-NUP153C and the CA resulted in the restriction of viral replication and a ∼2-fold drop in GFP positive cells, to 53%, compared to no TRIM control (Figure 3B). Of the eight NUP153C mutants that were predicted not to affect CA interaction (low ΔΔ*G*_bind_ values), five displayed wild type-like CA interaction reducing HIV-1 transduction to 69% (SD ±11.27) (P1411M, S1412M, G1413M, V1414I, and F1415M) (Figure 3B). The remaining three mutations were active in HIV-1 restriction but only reduced transduction to 81.21% (SD ± 4.87) (T1416M, F1417Y, and G1418A), indicating a loss of CA interaction. Five of the eight disruptive mutations with high ΔΔ*G*_bind_ values reduced transduction to 87.36% (SD ± 6.74) (V1414W, F1415G, T1417R, F1417G, and G1418Y) (Figure 3B). The less disruptive mutations with high ΔΔ*G*_bind_ values demonstrated almost wild-type-like restriction of HIV (P1411Y, S1412P, and G1413W).

**Figure 3.**
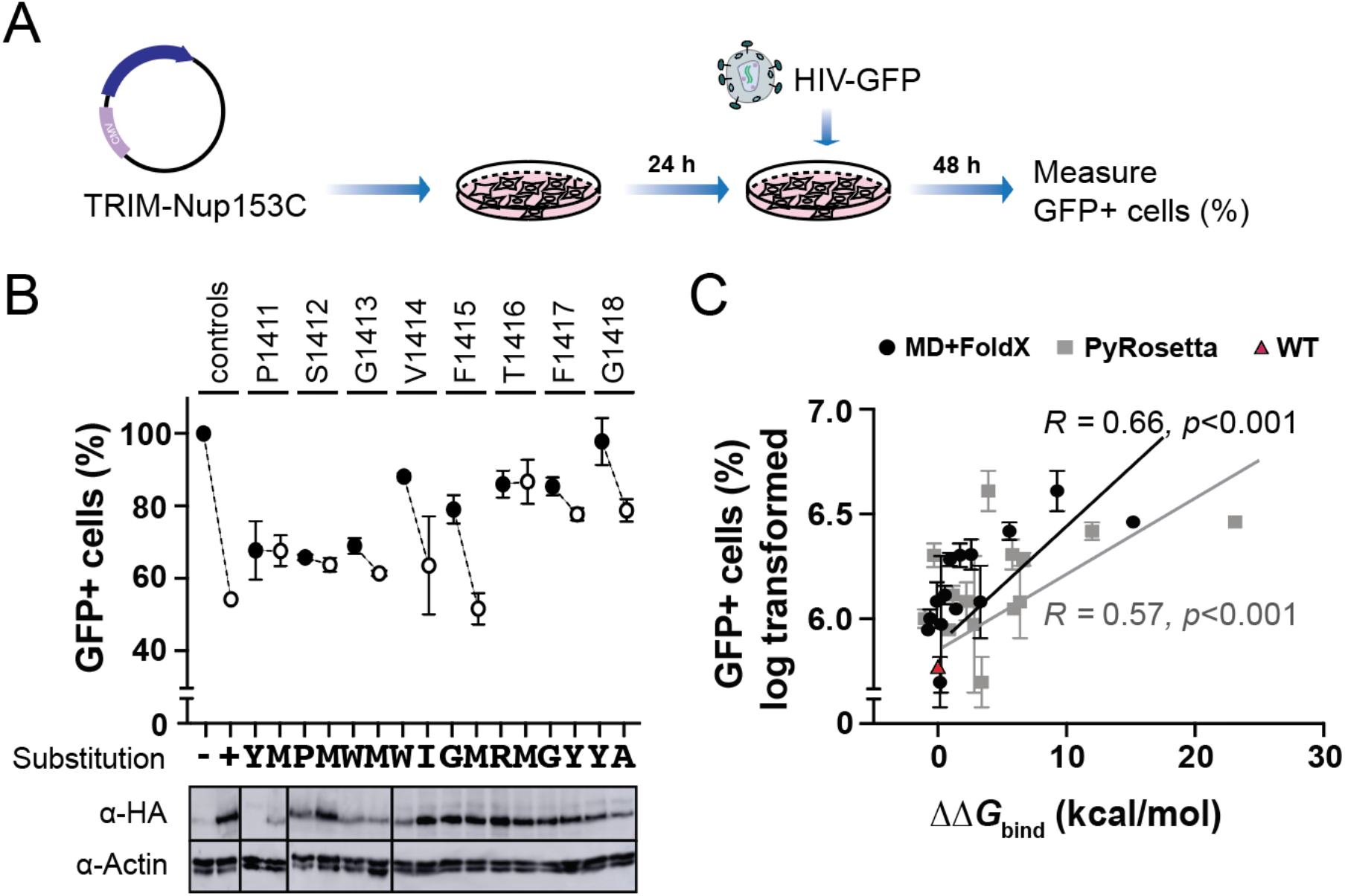
Molecular modeling predicts the effects of mutations in NUP153C on the CA interaction as measured by TRIM-NUP153C restriction. (A) A schematic representation of the workflow of a TRIM-NUP153C restriction assay. (B) *Top*. NUP153C interaction with capsid was measured by the degree of HIV-1 restriction by transient expression of TRIM-NUP153C in HEK293T cells. The relative percentage of GFP-positive cells, indicating HIV-1 transduction, was measured by flow cytometry. White data points represent mutations that had low ΔΔ*G*_bind_ values, and black circles represent mutations that had high ΔΔ*G*_bind_ values for each site. *Bottom*. The expression of each mutant TRIM-NUP153C was examined by Western blot. (C) The correlation of the empirical data presented in (B) with the ΔΔ*G*_bind_ as calculated by either MD+FoldX or PyRosetta.

Measuring the expression of the TRIM-NUP153C mutants indicated that the majority were expressed to a similar level within HEK293T cells (Figure 3B, bottom). When there was reduced expression of NUP153C, there did not appear to be a reduction in HIV restriction (P1411Y, P1411M, G1413W, G1413M). Of all the mutants tested, only V1414W showed decreased expression and a concomitant decrease in HIV restriction. As demonstrated with the co-sedimentation assay, we found a significant correlation between modeling predictions and TRIM-NUP153C restriction (Pearson’s correlation coefficient 0.66 (*p*<0.001) and 0.57 (*p*<0.001) for MD+FoldX and PyRosetta, respectively) (Figure 3C).

## Discussion

Using a multidisciplinary approach combining molecular modeling and empirical laboratory assays, we have identified the critical amino acid residues in the C-terminal domain of NUP153 required for CA interaction. We have determined that the minimal CA interaction motif is likely eight amino acids in length based on MD simulations and *in silico* mutagenesis using two different modeling software programs (MD+FoldX and PyRosetta). Specifically, we find that amino acids 1411-1418 (PSGVFTFG) remain stably associated with CA during MD simulations. Predictions made by both modeling approaches were in general agreement that the mutation of six of these eight residues would be disruptive to CA interaction. An *in vitro* co-sedimentation assay and a cell-based TRIM-NUP153C restriction assay were used to determine the biological relevance of the modeling predictions. Overall, MD+FoldX and PyRosetta predictions correlate well with empirical data, confirming the specific binding of CA to NUP153C. These data suggest that the minimal required sequence for the binding of NUP153C to CA is PxxVFTFG, where x is any amino acid.

Validation of MD+FoldX ΔΔ*G*_bind_ predictions using PyRosetta highlights a strong agreement between two different tools using different energy functions. MD+FoldX seemed to discern CA-interacting from non-interacting mutants better than PyRosetta, as confirmed by experimental data (Figures 2E and 3C). This difference between the models is likely because PyRosetta uses a single experimental structure of the NUP153-CA complex to predict ΔΔ*G*_bind_ values. In contrast, the MD+FoldX approach uses multiple snapshots extracted from MD simulations. Improved performance of MD+FoldX in predicting ΔΔ*G*_bind_ values also highlighted the importance of incorporating conformational sampling. Interestingly, both methods did poorly in predicting the effects of mutating T1416. Indeed, both methods suggest small ΔΔ*G*_bind_ values for T1416M and T1416R, indicating binding stabilization.

Conversely, experimental data has shown that the mutations T1416M and T1416R (this study) and T1416A (21) disrupt CA interaction. This discrepancy between modeling and empirical data is likely due to the water-mediated interaction of T1416 with CA residues R173 and E63 (21). These modeling inaccuracies are expected because one of the significant limitations of fast protein-protein binding affinity prediction tools (FoldX and PyRosetta) is that they ignore the explicit presence of bridging water molecules (31).

The hydrophobic pocket in the CA hexamer has been reported to accommodate the host factors CPSF6 and SEC24C (21–23). The same pocket has also been targeted by small molecules such as PF74 and BI-2 (21, 24), and the antiviral drug Lenacapavir and its derivatives (25, 32). These different proteins and small molecules adopt slightly different conformations within the CA pocket, but with a common feature of a phenyl group insertion into the binding pocket with hydrophobic interactions with CA. The phenyl group required to interact NUP153 with CA is the sidechain of F1417, which overlays with CA-bound CPSF6 residue F321 and Sec24C residue F236. In addition, the backbone amides of F1417, F321, and F236 all form a hydrogen bond with the sidechain of CA N57. However, bulky sidechain substitutions appear to be tolerated at F1417, as demonstrated by a significant correlation between increasing sidechain size and CA interaction (Table 1). Mutations at position F1417 were predicted to be deleterious to CA interaction with the lowest ΔΔ*G*_bind_ of 0.94 kcal/mol for the conservative mutation F1417Y. The average ΔΔ*G*_bind_ value of all mutations at F1417 was 3.71 kcal/mol (SD ± 1.44), demonstrating that F1417 is crucial for CA interaction. The importance of F1417 is confirmed by the empirical findings, except that the F1417Y substitution improved CA binding. Similarly, modeling predicted that mutation of F1415 was also deleterious to CA interaction, but the average ΔΔ*G*_bind_ value (1.85 kcal/mol; SD ± 0.78) was lower than that of F1417. Similar to F1417Y, the bulky F1415M mutation increased CA binding in both empirical assays and would be predicted to make hydrophobic contact with P38 of CA. The sulfur-containing sidechain of F1415M into this hydrophobic binding pocket of CA is similarly positioned to the sulfonyl group of Lenacapavir that hydrogen bonds with CA S41 and N57 (33). Both S1412 and G1413 were anchored to CA during MD simulations and were insensitive to mutations based on modeling predictions, and this agreed with empirical data. The ability of S1412 to tolerate mutation reflects the interaction with CA residue Q176 via the main chain. Similarly, G1413 can be substituted for tryptophan (G1413W) without altering CA interaction because the bulky sidechain would be exposed to the solvent. Even though these larger amino acids are accommodated at these positions, there is still a preference for small volume sidechains (Table 1).

Genetic variation in host factors hijacked by viruses can protect from viral infection and result in signatures of positive selection and NUP153 is evolving rapidly in primates (34). However, the CA interaction site is 100% identical in 35 primate species except in gorilla (*Gorilla gorilla*) and drill (*Mandrillus leucophaeus*). The fixed S1412P substitution in gorilla NUP153 would likely have a minor effect on CA interaction (ΔΔ*G*_bind_ 1.41 kcal/mol). Drill NUP153 has a more deleterious G1418S substitution (ΔΔ*G*_bind_ 2.27 kcal/mol). This substitution has a polar sidechain that is less likely to be accommodated within the hydrophobic CA pocket and limits the flexibility of the polypeptide backbone. Interestingly, a simian-human immunodeficiency virus (SHIV) chimera with the CA from SIV_mnd1_ (infecting *Mandrillus sphinx*) does not require NUP153 when infecting human cells (18). It is tempting to speculate that the incompatibility between SIV_mnd1_ and human NUP153 could be due to the adaptation of the SIV_mnd1_ CA to accommodate S1418 in this primate NUP153 or to circumnavigate NUP153 entirely. In humans, there were no high-frequency single nucleotide polymorphisms (SNPs) that would alter the amino acid sequence of the CA interaction site in NUP153 (gnomAD database (35). Two low-frequency non-synonymous SNPs were found at the same positions as in gorilla and drill resulting in the mutations S1412A and G1418V. These mutations in these individuals could potentially disrupt CA interaction and provide protection from HIV infection (ΔΔ*G*_bind_ 0.09 and 3.29 kcal/mol, respectively). Despite ongoing pressure from primate lentiviruses, there may have been selection against non-synonymous mutations at the CA interacting FG-containing motif of NUP153.

The minimal PxxVFTFG sequence required for CA interaction could mean that other FG repeats within NUP153 and other nucleoporins could bind CA. This is supported by the fact that only a complete deletion of the FG-region of NUP153 will abolish CA binding (20) and non-synonymous substitutions and small deletions do not completely perturb CA interaction (16). The relevance of other FG repeats in NUP153 for CA interaction remains to be thoroughly investigated.

## Materials and Methods

### Structure preparation for molecular modeling

The X-ray crystal structure of the HIV-1 CA hexamer interacting with human NUP153C was downloaded from Protein Data Bank (PDB ID:4U0D) (21). 3D coordinates file was modified to remove all but six chains of CA monomer and six chains of NUP153C. MODELLER software altered engineered residues in CA protein to wild-type and built the missing residues in all the chains to complete the experimental structure (36).

### Molecular dynamics simulations

The complete structure of the HIV CA hexamer bound to NUP153C was used as a starting structure for the MD simulation. The input structure was subjected to MD simulation using the protocol reported in our previous study (37). Briefly, the AMBER99SB*-ILDNP force field and the GROMACS 5.1.2 software package were used for generating topology files and performing simulations (38, 39). The final production simulation was run for 100 ns, and snapshots were saved every 1 ns resulting in 100 snapshots for the protein complex. The MD trajectory was visualized using the VMD software package and analyzed using the *grmsf* module available in the GROMACS package to calculate the root mean square fluctuation (RMSF) of all the atoms in each residue in the NUP153 motif during the simulation (40).

### Mutagenesis of NUP153C by FoldX and PyRosetta

MD snapshots of the HIV hexamer CA – NUP153 complex were analyzed using the FoldX software to estimate the relative binding affinities (ΔΔ*G*_bind_) for all possible mutations at each site in the NUP153 motif. As with our previous studies (27, 37), our FoldX analysis protocol involved processing each snapshot six times in succession using the RepairPDB command to energy minimize the snapshot and the BuildModel command to generate all possible 19 single mutations at each site in the NUP153 motif. The binding affinity (Δ*G*_bind_) was subsequently estimated using the AnalyseComplex command. ΔΔ*G*_bind_ for each mutation was calculated by taking the difference between mutated and wild-type Δ*G*_bind_ values. We then averaged ΔΔ*G*_bind_ values across all individual snapshot estimates for each mutation. To estimate ΔΔ*G*_bind_ values for all possible 19 mutations at each amino acid site of NUP153, we performed 30,400 FoldX calculations (16 NUP153 residues × 19 possible mutations at each site × 100 MD snapshots). Finally, we obtained 304 averaged ΔΔ*G*_bind_ values for all possible mutations of the NUP153 motif (<u>see S1 File</u>). PyRosetta-4 was used to compute the difference in binding stability scores between 16 selected mutant and wild-type structures (PDB: 4U0D) (41). The score is designed to capture the change in thermodynamic binding stability caused by the mutation (42). First, we repacked all sidechains sampled from the 2010 Dunbrack rotamer library (43) and applied the quasi-Newton minimization method via the ‘dfpmin’ algorithm in PyRosetta (44) with a tolerance of 0.001 (45) and the REF2015 scoring function (46) and allowing both the backbone torsion and sidechain angles to move. This procedure was performed ten times, and the lowest-scoring structure was selected for introducing mutations and subsequent binding stability calculations. Next, each missense mutation was introduced into the model of NUP153. All residues within a 10 Å distance of the mutated residue’s center were repacked, followed by a Monte Carlo sampling coupled with a quasi-Newton minimization of the backbone and all sidechains. We performed ten independent simulations of 5,000 Monte Carlo cycles each. To compute binding energy, we first scored the total energy of a bound state structure, separated CA and NUP153C, and then scored the unbound state total energy. The binding energy (Δ*G*_bind_) is computed by subtracting the unbound state total energy from the bound state total energy. This procedure was performed ten times, and the predicted ΔΔ*G*_bind_ was obtained by taking the average of the three lowest scoring structures. All molecular modeling data can be found in File S1.

### Plasmids construction and mutagenesis

The plasmid pLPCX-TRIM-NUP153C(human)-HA encoding the TRIM domain from TRIM5α of *Rhesus macaque* (residues 1 to 304) fused to the HA-tagged human NUP153 C-terminal domain (896 to 1475) was obtained from the Engelman laboratory. TRIM-NUP153C-HA was amplified by PCR and sub-cloned to the Gateway™ entry vector pCR8 to create plasmid pUI034. Gateway™ cloning introduced the gene into the custom destination vector pCDNA3-GW, following the manufacturer’s instructions (Thermo Fisher). Site-directed mutagenesis was performed using the Q5^®^ Site-Directed Mutagenesis Kit, following the manufacturer’s instructions (New England Biolab). All primers used for cloning and site-directed mutagenesis can be found in <u>Table S1</u> and a list of plasmids used in <u>Table S2</u>.

### Cells and virus

HEK293T cells (ACS-4500™, ATCC) were maintained at 37°C with 5% CO_2_ in Dulbecco’s Modified Eagle Medium (Sigma-Aldrich #D6429) supplied with 10% Fetal Bovine Serum (Sigma-Aldrich), 2 mM L-Glutamine (VWR #L0131-0100), and 1% Penicillin/Streptomycin solution (Corning, #30–002). Single-cycle HIV-1 virus with a GFP reporter gene was generated using a 10 mm dish by co-transfecting HEK293T cells with 4 μg psPAX2 (Addgene #12259), 4 μg pLJM1-EGFP (Addgene #19319), and 4 μg pCMV-VSVG (Addgene #8454) using Lipofectamine 3000 following the manufacturer’s instruction (Invitrogen). After 48 h, the supernatant was passed through a 0.45 μm filter and stored at - 80°C.

### Purification and in vitro assembly of CA hexamers and tubes

The expression of HIV-1 CA protein was adapted from Pornillos et al. (47). *E. coli* BL21(DE3)pLysS was transformed with pET11a-HIV-NL4-3 encoding CA with the mutations A14C and E45C. Transformed bacteria were cultured in LB media with ampicillin (15 μg/mL) and chloramphenicol (100 μg/mL) at 37°C until OD_600_ 0.8. Expression of CA was induced with a final concentration of 1 mM IPTG and incubated at 37°C for 4 h. Cells were centrifuged at 4,500 *x g* for 20 min at 4 °C. Cells were suspended in lysis buffer (50 mL for 4L; 50 mM Tris-Cl pH 8.0, 50 mM NaCl, 100 mM β-ME, protease inhibitor cocktail tablets [Sigma-Aldrich #11836153001]). The cell suspension was incubated on ice for 20 min with the addition of 1 g of lysozyme and 50 U of Benzonase® (EMD Millipore #70746-3). The cell suspension was subjected to sonication (MICROSON™ XL Ultrasonic cell disruptor) for 10 s at 80% of maximum output power for a total processing time of 5 min. Between pulses, samples were allowed to cool for 30 s on ice. The cell lysate was clarified by centrifugation (27,000 *x g* for 1 h at 4°C) and incubated with supersaturated ammonium sulfate (final concentration 25% of the total volume) on ice for 20 min. Precipitated CA was collected by centrifugation at 9,000 *x g* for 20 min at 4°C. The pellet was suspended in dialysis buffer (20 mM MOPS pH 6.8, 20 mM β-ME), transferred to a 3.5K MWCO dialysis cassette (Thermo Fisher #PI66110), and dialyzed against 1 L dialysis buffer for 16 h.

CA was purified using ion-exchange chromatography (ÄKTA start protein purification system, Cytiva). Dialysed lysate was centrifuged at 20,000 *x* g for 10 min at 4°C, passed through a 0.45 μm filter, and applied to a HiTrap SP FF (Cytiva #17-5054-01) column connected to a HiTrap Q FF column (Cytiva #17-5156-01). Fractions were collected at ∼25% of the sodium chloride gradient (∼0.25 M) and assayed for purity by SDS-PAGE (Fig S1A). Eluted CA protein was dialyzed using a 10K MWCO cassette (Thermo Fisher #PI66130). To assemble CA oligomers, the cassette was sequentially incubated at 4°C in assembly buffer (25 mM Tris-Cl pH 8.0, 1M NaCl) supplemented with 20 mM, 2 mM, and 0.2 mM β-ME for 8 h, 24 h, and 48 h, respectively. The efficiency of CA assembly into hexamers was assessed by SDS-PAGE, and assembly into multimeric tubes was confirmed by transmission electron microscopy (Franceschi Microscopy & Imaging Center, Washington State University).

### CA co-sedimentation assay

This assay was adapted from Selyutina et al. (48). Approximately 60,000 HEK293T cells were seeded into each well of a 12-well dish. After incubation for 24 h, cells were transfected with 500 ng of pCDNA3-TRIM-NUP153C using 1.5 μL of TransIT®-293 transfection reagent (Mirus Bio). Cells were incubated for 24 h before harvesting by scraping into 100 μL of CA binding buffer (10 mM Tris, pH 7.4, 1.5 mM MgCl_2_, 10 mM KCl, 1X Halt™ protease and phosphatase inhibitor cocktail [Thermo Fisher #PI78440]). Cell lysates were mixed for 15 min at 4°C before centrifugation at 21,000 *x g* for 15 min at 4°C. Clarified cell lysates were collected, and the protein content was normalized to 1.5 mg/mL by Bradford assay. 20 μL of CA tubes (∼112 pmol) and 80 μL of whole-cell lysate were mixed and incubated at room temperature for 1 h. The reaction was centrifuged for 8 min at 21,000 *x g* at 4°C. 15 μL of the supernatant was collected and compared to samples that were not centrifuged using Western dot blotting.

### Western blotting

Samples were separated by SDS-PAGE and transferred to nitrocellulose membranes by Trans-Blot® Turbo™ Transfer System (1.0 A, 25V, 15 min). Alternatively, samples were loaded onto the nitrocellulose membrane using a Bio-Dot® microfiltration apparatus (Bio-Rad). Membranes were blocked with 3% non-fat milk in TBS with 0.1% Tween-20 (TBST) for 1 h. To probe for the HA tag, the membrane was incubated with rat anti-HA-HRP (3F10, Sigma-Aldrich #12013819001; 1 in 2,000 dilution) for 1 h. To probe for CA and actin, membranes were incubated with rat anti-p24 antibody (ARP-64571; NIH HIV Reagent Program; 1 in 5,000 dilution) or rat anti-actin antibody (clone C4, VWR #10221-880; 1 in 500 dilution) for 1 h. Membranes were washed with 5 mL of TBST three times for 5 min each with gentle rocking. The anti-p24 blots were transferred to a new tray and probed against goat anti-rat antibody (Thermo Fisher #62-652-0; 1 in 4,000 dilution) for 40 min. Blots were visualized, and signals were quantified using Amersham™ Imager 600. Exposure time was adjusted manually to ∼10 seconds for anti-HA blots, ∼4 s for anti-p24 blots, and ∼10 seconds for anti-actin blots.

### TRIM-NUP153C-mediated restriction and cell flow cytometry

This assay was adapted from Matreyek et al. (16). 60,000 HEK293T cells were seeded and transfected with TRIM-NUP153C as described in the CA co-sedimentation assay. 24 h post-transfection, cells were transduced with HIV-GFP. Media was discarded 24 h post-transduction, and fresh media was added to the wells. 48 h after transduction, cells were treated with 0.25% trypsin (VWR #16777-202) and centrifuged at 2,000 *x g* for 3 min at room temperature. Cell pellets were suspended, fixed with 300 μL of Dulbecco’s phosphate-buffered saline (DPBS; VWR #45000-434) containing 1% paraformaldehyde (Electron Microscopy Sciences #15710), and incubated at 4°C for 1 h. Cells were centrifuged at 2,000 *x g* for 3 min, and the cell pellet was suspended in 500 μL DPBS. This step was repeated twice, and the final pellet was suspended in 100 μL flow cytometry buffer (DPBS with 4% FBS) and transferred to a 96-well U bottom assay plate (CELLTREAT #229590). Fluorescent cells were quantified using the CytoFLEX S instrument (Beckman Coulter).

## Author Contributions

S.L., J.S.P., J.Y., and P.A.R. wrote the manuscript.

## Acknowledgment

We would like to thank Dr. Lee Fortunato for the advice and technical support in creating HIV-GFP. We also thank Yesol Sapozhnikov, Dr. Craig Miller, and Dr. Mohamed Megheib for their help with statistical analyses. The following reagent was obtained through the NIH HIV Reagent Program, Division of AIDS, NIAID, NIH: Monoclonal Anti-Human Immunodeficiency Virus Type 1 (HIV-1) p24 Gag (#24-2, produced *in vitro*), ARP-6457, contributed by Dr. Michael H. Malim.

